# Drug Repurposing Screen for Compounds Inhibiting the Cytopathic Effect of SARS-CoV-2

**DOI:** 10.1101/2020.08.18.255877

**Authors:** Catherine Z. Chen, Paul Shinn, Zina Itkin, Richard T. Eastman, Robert Bostwick, Lynn Rasmussen, Ruili Huang, Min Shen, Xin Hu, Kelli M. Wilson, Brianna Brooks, Hui Guo, Tongan Zhao, Carleen Klump-Thomas, Anton Simeonov, Samuel G. Michael, Donald C. Lo, Matthew D. Hall, Wei Zheng

**Author notes:** To whom correspondence should be addressed: Catherine Chen –, Wei Zheng –.

## Abstract

Drug repurposing is a rapid approach to identifying therapeutics for the treatment of emerging infectious diseases such as COVID-19. To address the urgent need for treatment options, we carried out a quantitative high-throughput screen using a SARS-CoV-2 cytopathic assay with a compound collection of 8,810 approved and investigational drugs, mechanism-based bioactive compounds, and natural products. Three hundred and nineteen compounds with anti-SARS-CoV-2 activities were identified and confirmed, including 91 approved drug and 49 investigational drugs. Among these confirmed compounds, the anti-SARS-CoV-2 activities of 230 compounds, including 38 approved drugs, have not been previously reported. Chlorprothixene, methotrimeprazine, and piperacetazine were the three most potent FDA approved drugs with anti-SARS-CoV-2 activities. These three compounds have not been previously reported to have anti-SARS-CoV-2 activities, although their antiviral activities against SARS-CoV and Ebola virus have been reported. These results demonstrate that this comprehensive data set of drug repurposing screen for SARS-CoV-2 is useful for drug repurposing efforts including design of new drug combinations for clinical trials.

## Introduction

The coronavirus disease 2019 (COVID-19) pandemic caused by severe acute respiratory syndrome coronavirus 2 (SARS-CoV-2) has become a global health crisis. As of July 31, 2020, the global case report stands at 17.5 million, with a death toll of 677,538 (Dong et al., 2020). Only remdesivir, an investigational drug developed for Ebola virus, has been granted emergency use authorization by the US FDA (Eastman et al., 2020). Since neither an effective vaccine, nor antiviral therapy, are currently available for COVID-19, drug repurposing has received significant attention in the rapid search to fill this unmet therapeutic need.

The requirement of biosafety level 3 (BSL-3) containment laboratories for handling SARS-CoV-2 has limited the number of high throughput screening (HTS) laboratories that are capable of carrying out large scale compound screens using live SARS-CoV-2. Despite these challenges, several drug repurposing screens have been carried out using live SARS-CoV-2, showing promising results (Dittmar et al., 2020; Ellinger et al., 2020; Riva et al., 2020; Touret et al., 2020). Here we report a screening campaign against a collection of 8,810 approved and investigational drugs, mechanism-based bioactive compounds, and natural products that was carried out in quantitative HTS (qHTS) format (Inglese et al., 2006). Compounds were screened at four concentrations in a SARS-CoV-2 cytopathic effect (CPE) assay in Vero E6 cells that were selected for high ACE2 expression, with an accompanying cytotoxicity counter-assay. The primary screen yielded 319 hits with confirmed anti-SARS-CoV-2 activity. The primary screening data have been made publicly available on the NCATS OpenData Portal (https://opendata.ncats.nih.gov/covid19/index.html) (Brimacombe et al., 2020). We intend this as a companion to guide investigators in utilizing that data, and to present further details of qHTS with the SARS-CoV-2 CPE assay, including identification of top annotated hits.

## Materials and Methods

### Compounds and compound libraries

All compound libraries were assembled internally at NCATS. The NCATS pharmaceutical collection (NPC) contains 2,678 compounds, covering drugs approved by US FDA and foreign health agencies in European Union, United Kingdom, Japan, Canada, and Australia, as well as some clinical trialed experimental drugs (Huang et al., 2019). The NCATS Mechanism Interrogation Plate (MIPE) 5.0 library contains 2,480 mechanism based bioactive compounds, targeting more than 860 distinct mechanisms of action (Lin et al., 2019). The NCATS Pharmacologically Active Chemical Toolbox (NPACT) is a library of mechanistically defined molecules and natural products (5,099 compounds). Other small custom NCATS collections were also screened: anti-infective (752 compounds), kinase inhibitors (977 compounds), epigenetic modulators (335 compounds). A commercially available autophagy-focused screening library (Cayman #23537) was analyzed and 29 compounds that were not already present in our collections were purchased. All compounds were dissolved in DMSO to make 10 mM stock solutions, unless solubility was limiting, and was diluted four times at 1:5 ratio for the primary screens, and at 1:3 ratio for follow up assays at 8 concentrations.

### CPE assay

A SARS-CoV-2 CPE assay was conducted in the BSL3 facilities at the contract research organization Southern Research (Birmingham, AL). Briefly, compounds were titrated in DMSO and acoustically dispensed into 384-well assay plates at 60 nL/well at NCATS, and provided to Southern Research. Cell culture media (MEM, 1% Pen/Strep/GlutaMax, 1% HEPES, 2% HI FBS) was dispensed at 5 μL/well into assay plates and incubated at room temperature to allow for compound dissolution. Vero E6 African green monkey kidney epithelial cells (selected for high ACE2 expression) were inoculated with SARS CoV-2 (USA_WA1/2020) at multiplicity of infection (MOI) of 0.002 in media and quickly dispensed into assay plates as 25 μL/well. The final cell density was 4000 cells/well. Assay plates were incubated for 72 h at 37C, 5% CO_2_, 90% humidity. CellTiter-Glo (30 μL/well, Promega #G7573) was dispensed into the assay plate. Plates were incubated for 10 min at room temperature. Luminescence signal was measured on Perkin Elmer Envision or BMG CLARIOstar plate readers. An ATP content cytotoxicity counter-assay was conducted using the same protocol as the CPE assay, without the addition of SARS-CoV-2 virus.

### Data analysis

Results from the primary screen and confirmation screens were processed at NCATS using a software developed in-house (Wang et al., 2010). For the CPE assay, raw plate data were normalized with DMSO-only wells as 0% CPE rescue (negative signal control) and no virus control wells as 100% CPE rescue (positive signal control). For the cytotoxicity assay, raw plate data were normalized with DMSO-only wells as 100% viability (positive signal control) and cells treated with hyamine (benzethonium chloride) control compound as 0% viability (negative signal control). The half-maximum effective values (EC_50_) and percent efficacy were obtained by fitting the concentration-response titration data to a four-parameter Hill equation. Compounds with >55% efficacy were selected for cherry-pick confirmation. The concentration-response curves of re-tested compounds were also plotted using GraphPad Prism 9 (GraphPad Software Inc., San Diego, CA). Results in the figures are expressed as mean ± standard deviation (SD).

## Results

### High throughput screening with SARS-CoV-2 CPE assay

Our aims were two-fold in initiating this program. The first was to identify active compounds that may provide opportunities for repurposing or identify mechanistic targets of interest. The second was to create a complete HTS reference dataset that can be shared openly with the scientific community. The CPE reduction assay format has been widely employed to screen for antiviral agents due to its easy of scalability for HTS (Heaton, 2017). In this assay, viral infection kills host cells and the cell viability is used as a surrogate readout for viral infection and replication. In other words, compounds with anti-viral activity rescue cells from the cytopathic effect of SARS-CoV-2 (a gain-of-signal assay).

A total of 9,952 compounds were tested in the primary screen, but due to overlap in the composition of libraries, a significant number of compounds were tested multiply. A total of 8,810 unique compounds in six compound libraries were tested in the primary screen including the NCATS Pharmaceutical Collection (NPC), NCATS Mechanism Interrogation Plate (MIPE), NCATS Pharmacologically Active Chemical Toolbox (NPACT), Epigenomic library, Autophagy library, and anti-infective library. These compounds contain 1,345 approved drugs (by FDA, EMA, DPD), 751 compounds approved outside of these countries, 1067 investigational drugs (in clinical trials), 1057 pre-clinical compounds (tested in animals) and 4,472 bioactive compounds (tool compounds) (Figure 1a). By their mechanisms of action and clinical application, these compounds are divided into diverse groups (Figure 1b).

**Figure 1.**
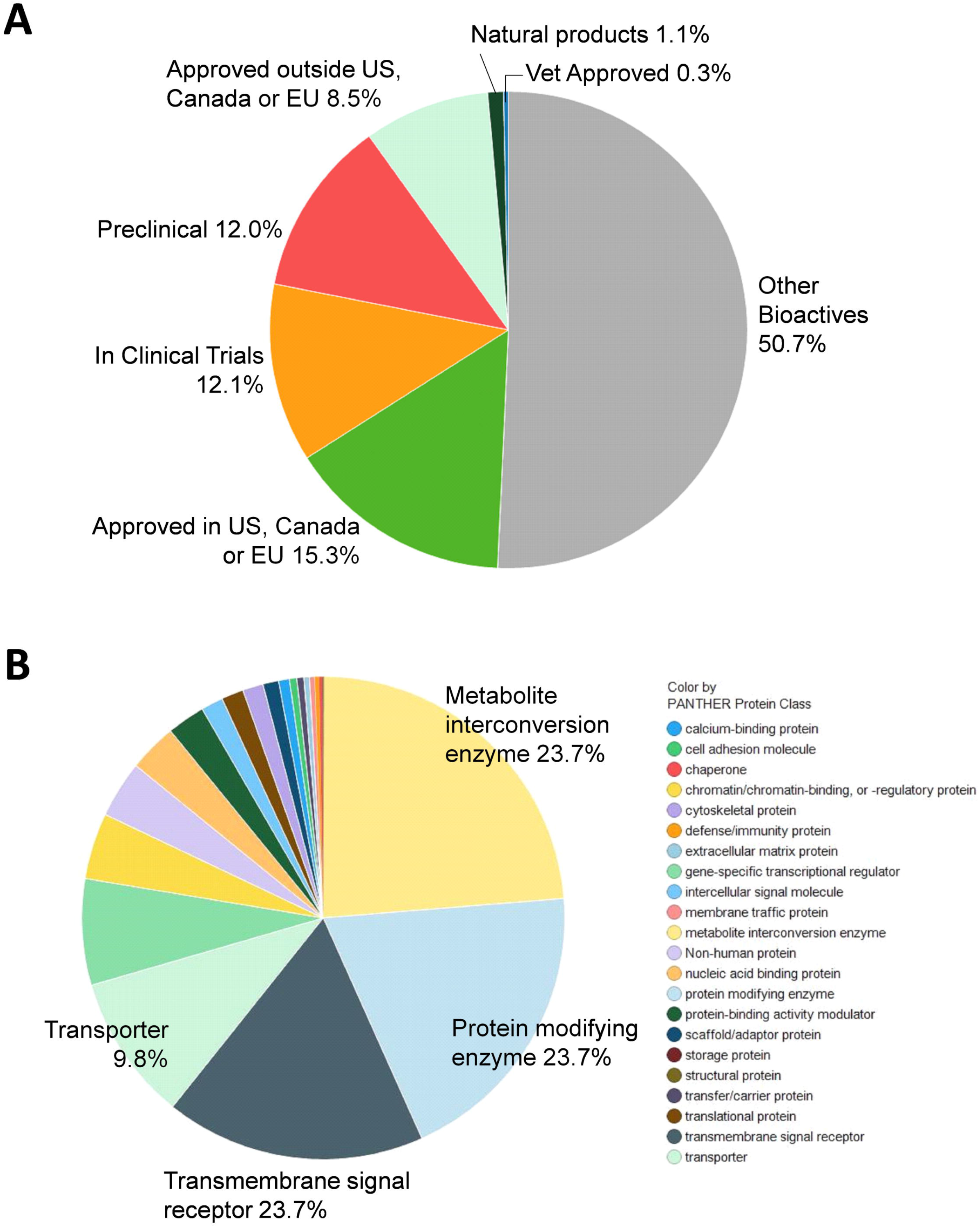
Compound library description. (a) By approval status: approved drug (FDA and others), clinical trial, preclinical. (b) By mechanism of action.

The CPE assay performed well in the primary screen, with an average Z’ factor of 0.83 over 133 plates over three batched runs (Figure 2a). Remdesivir concentration response was included as a control for each screening run and yielded consistent EC50 values of 4.56 μM, 4.42 μM and 7.28 μM (Figure 2b). Using the criteria of >55% efficacy, 380 compounds were selected as the primary screen hits, out of which, 319 compounds were confirmed using 8-point, 1:3 titration in duplicate. Among these primary hits, 89 of 319 had previously reported activity against SARS-CoV-2, including the assays of live virus, enzymatic, or virtual screening, while 230 were novel hits from this qHTS (Table 1, S1).

**Figure 2.**
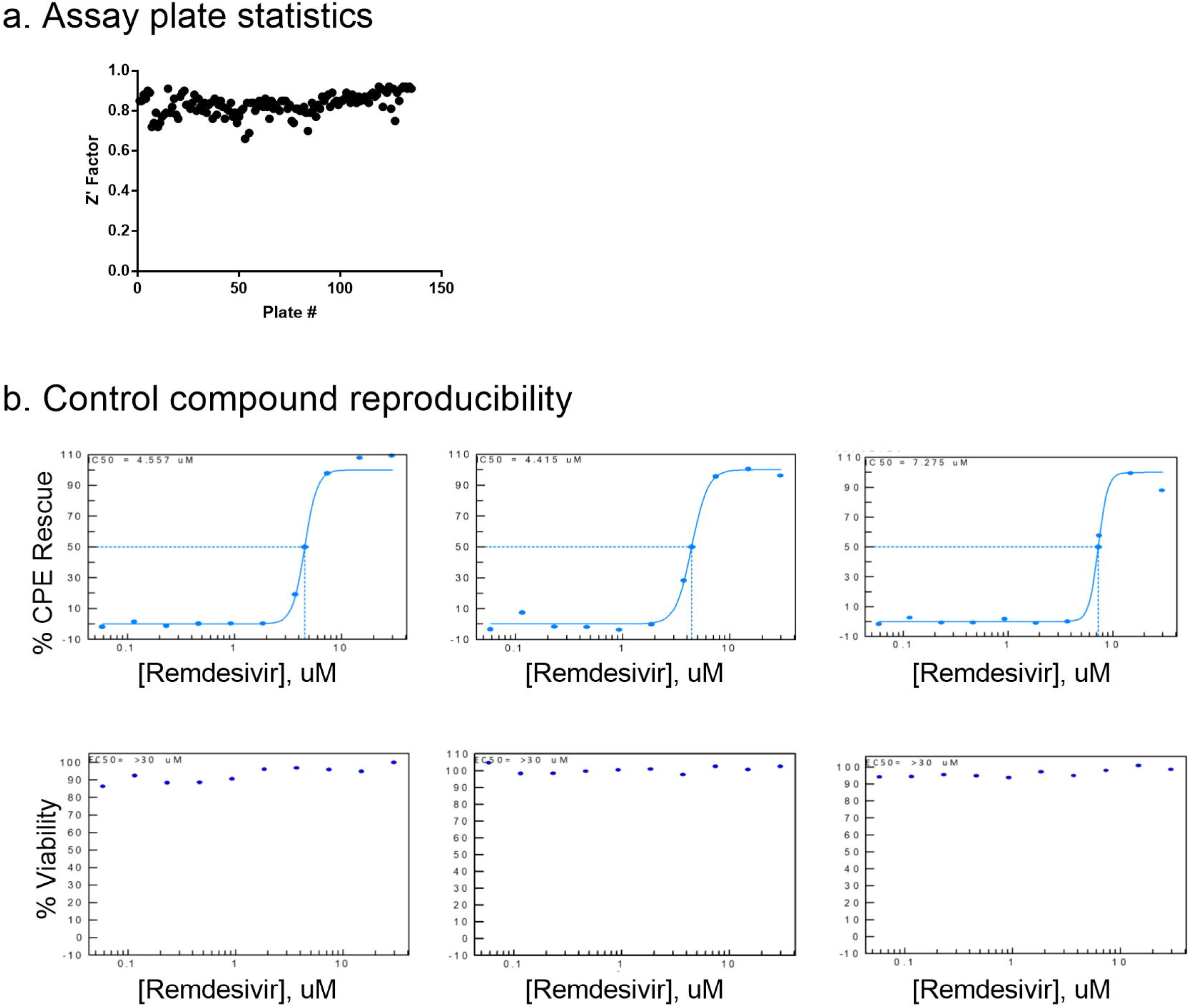
Control compounds reproducibility. Concentration response curve fittings for remdesivir in four independent runs for primary screens and hit confirmation. EC50 values of 4.56 μM, 4.42 μM, 7.28 μM, and 5.17 μM of remdesivir in the CPE assay demonstrate day-to-day reproducibility of the assay.

**Table 1.** Top confirmed anti-SARS-CoV-2 compounds.

| Sample ID | Sample Name | CPE EC50 (uM) | CPE % Efficacy | Cytotox CC50 (uM) | % Cytotox | Previous reports against CoVs | Approval status | MOA |
| --- | --- | --- | --- | --- | --- | --- | --- | --- |
| **Viral Target** |  |  |  |  |  |  |  |  |
| NCGC00686694 | Remdesivir | 10.0 | 133.1 | N/A | <30 | Clinical (Beigel et al., 2020) | FDA EUA | RdRP inhibitor |
| **Autophagy Modulators** |  |  |  |  |  |  |  |  |
| NCGC00387732 | VPS34-IN1 | 0.63 | 103.0 | 10.0 | -76.5 | None | Bioactive | Autophagy modulator |
| NCGC00344081 | STF-62247 | 1.1 | 107.1 | 11.2 | -56.6 | None | Preclinical | Autophagy modulator; Renal cell growth inhibition |
| NCGC00507892 | VPS34 Inhibitor 1 | 1.4 | 98.3 | N/A | <30 | None | Preclinical | Autophagy modulator |
| **GPCR Modulators** |  |  |  |  |  |  |  |  |
| NCGC00346896 | MCOPPB | 3.5 | 85.6 | N/A | <30 | None | Preclinical | ORL1 (OP4, NOP) Agonists |
| NCGC00370950 | GW 803430 | 3.5 | 93.3 | N/A | <30 | None | Bioactive | Melanin-concentrating hormone receptor 1 Antagonist |
| NCGC00017063 | Amodiaquin dihydrochloride | 4.0 | 87.2 | N/A | <30 | In vitro live virus (Weston et al., 2020) | FDA | Histamine receptor antagonist |
| NCGC00485045 | N-Methylspiperone hydrochloride | 4.5 | 80.0 | N/A | <30 | None | Clinical trial | Serotonin 2 (5-HT2) receptor Antagonist |
| NCGC00016710 | Clemastine fumarate | 7.9 | 96.0 | N/A | <30 | Mpro assay (Vatansever et al., 2020) | FDA | Histamine receptor antagonist |
| NCGC00386477 | GMC 2-29 | 7.9 | 117.2 | N/A | <30 | None | Bioactive | 5-hydroxytryptamine receptor 1D Antagonist |
| NCGC00378842 | Lu AE58054 hydrochloride | 10.0 | 97.2 | N/A | <30 | None | Clinical trial | Serotonin 6 (5-HT6) receptor Antagonist |
| NCGC00013683 | Chlorprothixene | 10.0 | 104.4 | N/A | <30 | None | FDA | Dopamine receptor antagonist |
| NCGC00014482 | Methdilazine hydrochloride | 10.0 | 86.4 | N/A | <30 | Virtual: AI prediction (Grzybowski et al., 2020) | FDA | Antihistamine |
| NCGC00179370 | Methotrimeprazine maleate | 10.0 | 84.6 | N/A | <30 | None | FDA | Antagonist for adrenergic, dopamine, histamine, cholinergic and serotonin (5-hydroxytryptamine; 5-HT) receptors |
| NCGC00016642 | Piperacetazine | 10.0 | 103.7 | N/A | <30 | None | FDA | Dopamine receptor antagonist |
| NCGC00181913 | Difeterol | 10.0 | 113.4 | N/A | <30 | None | Approved outside of US | Antihistamine |
| NCGC00386484 | (R)-(-)-LY 426965 dihydrochloride | 10.0 | 110.7 | N/A | <30 | None | Bioactive | Serotonin 2b (5-HT2b) receptor Modulator |
| NCGC00015608 | Loperamide hydrochloride | 10.0 | 98.6 | N/A | <30 | In vitro live virus (Jeon et al., 2020) | FDA | Opioid receptor agonist |
| NCGC00485321 | Naltrindole isothiocyanate hydrochloride | 10.0 | 114.7 | N/A | <30 | None | Bioactive | Delta opioid receptor Antagonist |
| NCGC00165726 | AM1241 | 10.0 | 97.6 | N/A | <30 | None | Bioactive | Cannabinoid CB2 receptor Agonist |
| NCGC00386703 | CpdD hydrochloride | 10.0 | 96.9 | N/A | <30 | None | Bioactive | Ghrelin receptor Antagonist |
| NCGC00386219 | SB 271046 hydrochloride | 10.0 | 107.5 | N/A | <30 | None | Bioactive | Serotonin 6 (5-HT6) receptor Antagonist |
| NCGC00386479 | GMC 2-113 | 10.0 | 129.7 | N/A | <30 | Virtual: RdRP (Dwivedy et al., 2020) | Bioactive | 5-hydroxytryptamine receptor 1D Antagonist |
| **Host Protease Inhibitors** |  |  |  |  |  |  |  |  |
| NCGC00386330 | Z-FA-FMK | 0.13 | 104.8 | N/A | <30 | Mpro assay, in vitro live virus (Zhu et al., 2020b) | Bioactive | Cathepsin L Inhibitor |
| NCGC00485951 | VBY-825 | 0.14 | 97.8 | N/A | <30 | In vitro live virus (Riva et al., 2020) | Clinical trial | Cathepsin S Inhibitor |
| NCGC00345807 | CAA-0225 | 0.20 | 99.3 | N/A | <30 | None | Preclinical | Cathepsin L Inhibitors |
| NCGC00386232 | Cathepsin Inhibitor 1 | 0.25 | 114.4 | N/A | <30 | None | Bioactive | Cathepsin inhibitors |
| NCGC00163432 | Calpeptin | 0.50 | 111.7 | N/A | <30 | Mpro assay, in vitro live virus (Ma et al., 2020) | Preclinical | Calpain inhibitor |
| NCGC00485375 | Z-Gly-Leu-Phe-chloromethyl ketone | 1.3 | 87.2 | N/A | <30 | None | Bioactive | Granzyme B Inhibitor |
| NCGC00371151 | Balicatib | 2.0 | 100.3 | N/A | <30 | None | Clinical trial | Cruzain (Trypanosoma cruzi) Inhibitor |
| NCGC0016166 | Calpain Inhibitor I, ALLN | 2.0 | 111.1 | N/A | <30 | None | Bioactive | Calpain inhibitor |
| **Kinase Modulators** |  |  |  |  |  |  |  |  |
| NCGC00263093 | Apilimod | 0.023 | 104.4 | N/A | <30 | Pseudovirus assay (Ou et al., 2020) | Clinical trial | IL-12 Production Inhibitor; PIKfyve inhibitor |
| NCGC00386313 | Berzosertib | 0.71 | 87.9 | 11.2 | -98.5 | None | Clinical trial | ATR Kinase Inhibitor |
| NCGC00347280 | IKK-2 Inhibitor VIII | 7.1 | 91.7 | N/A | <30 | None | Preclinical | IKK-2 (IKK-beta) Inhibitor |
| NCGC00387166 | NSC 33994 | 8.9 | 107.6 | N/A | <30 | None | Bioactive | Jak2 Inhibitor |
| NCGC00159456 | Imatinib | 10.0 | 119.0 | N/A | <30 | Clinical (Morales-Ortega et al., 2020) | FDA | Bcr-Abl kinase inhibitor; KIT inhibitor; PDGFR tyrosine kinase receptor inhibitor |
| **Others** |  |  |  |  |  |  |  |  |
| NCGC00178090 | Pristimerin | 0.11 | 87.4 | 1.1 | -93.2 | SARS Mpro assay (Ryu et al., 2010) | Preclinical | Monoacylglycerol lipase (MGL) inhibitor |
| NCGC00385252 | alpha-L-Arabinopyranose | 2.4 | 104.0 | N/A | <30 | None | Bioactive | Induces Pbad promoter expression in E. coli |
| NCGC00351072 | ML414 | 3.2 | 79.6 | N/A | <30 | None | Bioactive | Oligosaccharyltransferase inhibitor |
| NCGC00379165 | IT1t dihydrochloride | 3.5 | 96.3 | N/A | <30 | None | Bioactive | CXCR4 inhibitor |
| NCGC00485648 | S-15176 difumarate salt | 3.8 | 127.4 | N/A | <30 | None | Bioactive | Oxidative Stress Inhibitor |
| NCGC00384450 | JTV519 Hemifumarate | 5.5 | 85.7 | N/A | <30 | None | Clinical trial | Ryanodine receptor (RyR) inhibitor |
| NCGC00253604 | Rescimetol | 8.9 | 81.8 | N/A | <30 | None | Approved outside of US | Antihypertensive agent |
| NCGC00164559 | Duloxetine hydrochloride | 10.0 | 90.0 | N/A | <30 | Mpro assay (Vatansever et al., 2020) | FDA | Norepinephrine reuptake inhibitor; Serotonin-norepinephrine reuptake inhibitor (SNRI) |
| NCGC00181168 | Trifluomeprazine 2-butenedioate | 10.0 | 90.2 | N/A | <30 | None | Bioactive | Antipsychotic Agents |
| NCGC00169804 | Asteriscunolide D | 10.0 | 93.3 | N/A | <30 | None | Bioactive | Natural product |
| NCGC00485925 | Genz-123346 (free base) | 10.0 | 99.4 | N/A | <30 | In vitro live virus (Vitner et al., 2020) | Bioactive | Ceramide glucosyltransferase Inhibitor |
| NCGC00015708 | Maprotiline hydrochloride | 10.0 | 103.7 | N/A | <30 | Virtual: Mpro docking (Chauhan, 2020) | FDA | Norepinephrine reuptake inhibitor; Tricyclic antidepressant |
| NCGC00168786 | Deserpidine | 10.0 | 84.7 | N/A | <30 | Virtual: NSP16 docking (Jiang et al., 2020) | FDA | Angiotensin converting enzyme inhibitor |
| NCGC00015096 | Amiodarone hydrochloride | 10.0 | 100.5 | N/A | <30 | Clinical (Castaldo et al., 2020) | FDA | Potassium channel blocker |
| NCGC00181088 | Melitracen hydrochloride | 10.0 | 97.1 | N/A | <30 | None | Approved outside of US | Antidepressive Agents, Tricyclic; |
| NCGC00015428 | (+/-) - Fluoxetine | 10.0 | 115.8 | N/A | <30 | In vitro live virus (Zimniak et al., 2020) | FDA | Selective serotonin reuptake inhibitor (SSRI) |
| NCGC00018102 | Flunarizine | 10.0 | 94.1 | N/A | <30 | Virtual: Spike docking (Chernyshev, 2020) | Approved outside of US | Calcium channel blocker |
| NCGC00183024 | Proglumetacin | 10.0 | 87.6 | N/A | <30 | None | Approved outside of US | Cyclooxygenase inhibitor |
| NCGC00378760 | DMP 777 | 10.0 | 92.5 | N/A | <30 | None | Clinical trial | Leukocyte elastase Inhibitor |
| NCGC00476094 | Dexanabinol | 10.0 | 110.8 | N/A | <30 | None | Clinical trial | NMDA antagonist |

### 91 approved drugs and 49 investigational drugs protected against cytopathic effect of SARS-CoV-2 infection

There were 56 top confirmed hits with EC_50_ values of ≤10 μM and efficacy greater than 80% in the CPE assay, and greater than 10-fold selectivity index between cytotoxicity and CPE assays (Table 1, Figure 3). When grouped by mechanism of action targets, 19 compounds were GPCR modulators, 8 were host protease inhibitors, 5 were kinase modulators, and 3 were autophagy modulators (Figure 3). Interestingly, in the 56 top hits, only one, remdesivir has a known viral target as a known primary mechanism, whereas the known mechanisms of action of the other compounds are directed against host targets.

**Figure 3.**
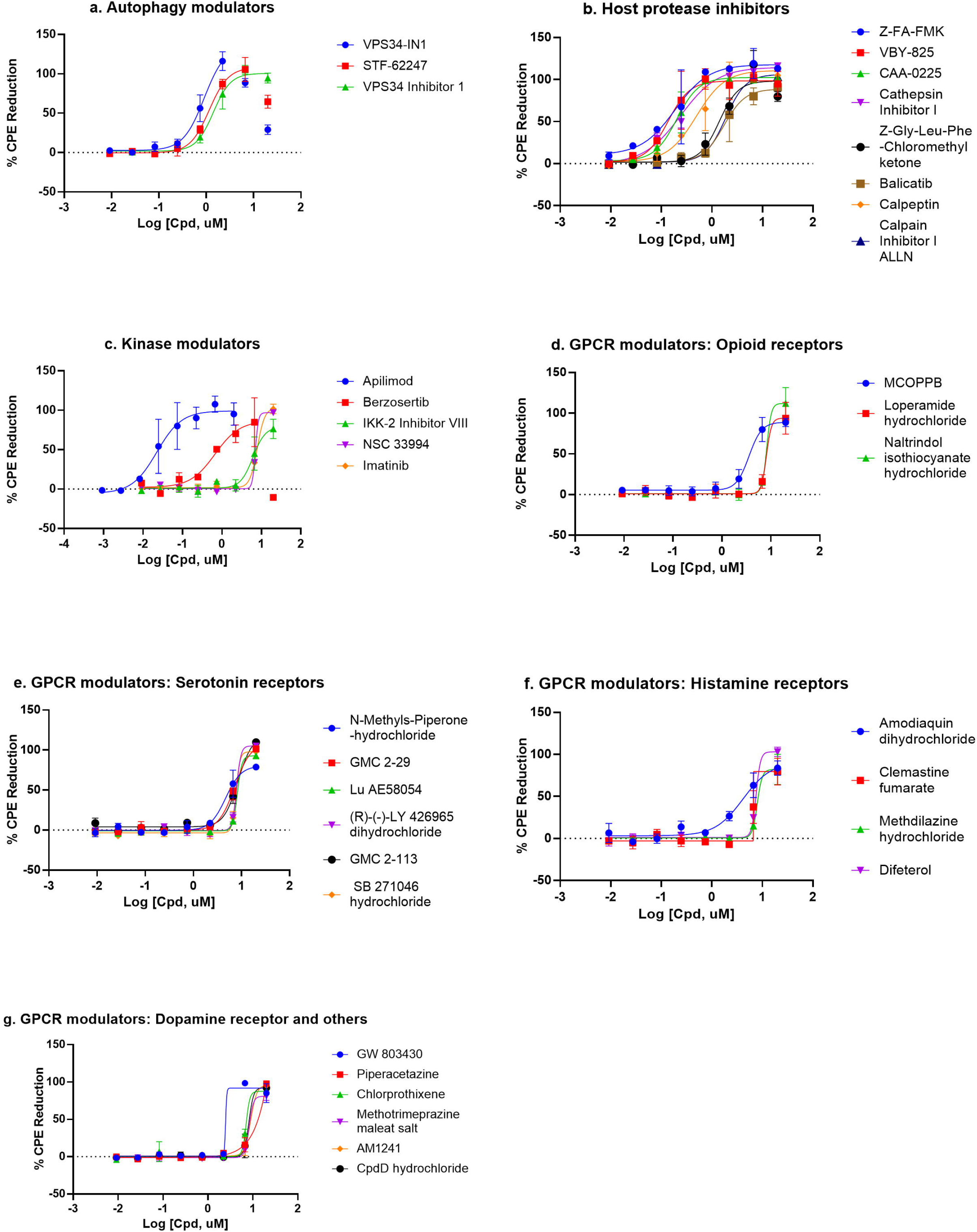
Compounds concentration response curves in the CPE assay. (a) Autophagy modulators, (b) host protease inhibitors, (c) kinase modulators, (d) opioid receptor modulators, (e) serotonin receptor modulators, (f) histamine receptor modulators, and (g) dopamine and other GPCR receptor modulators. Berzosertib, VPS34-IN1, and STF-62247 showed bell-shaped concentration response due to cytotoxicity. No other compounds caused any reduction in viability in the cytotoxicity assay.

There have been several previous drug repurposing screens reported for SARS-CoV-2 in 2D cell culture infection models (Dittmar et al., 2020; Ellinger et al., 2020; Jeon et al., 2020; Riva et al., 2020; Touret et al., 2020; Weston et al., 2020). These screens had some compound overlap with our qHTS screen, particularly for the FDA approved drugs. We performed a literature search of our confirmed compounds and previous reports were noted in Table 1 and Table S1. Three of the top 56 hits were novel and FDA approved. These hits are chlorprothixene, methotrimeprazine, and piperacetazine, which showed 10 μM potencies in the CPE assay. Four drugs approved outside of the US were also identified as novel compounds with anti-SARS-CoV-2 effects: difeterol, rescimetol, melitracen HCl, and proglumetacin. Furthermore, we identified 7 novel clinical trial drugs with anti-SARS-CoV-2 activities: N-methylspiperone HCl, Lu AE58054 HCl, balicatib, berzosertib, JTV519 hemifumarate, DMP 777, and dexanabinol. In addition to above novel hits, four drugs, approved by FDA and elsewhere, methdilazine, maprotiline HCl, deserpidine, and flunarizine, were previously reported in virtual screens against SARS-CoV-2 targets without supporting biological data. Here, we report their activities against SARS-CoV-2 infection. In addition, we have confirmed 53 approved drugs with anti-SARS-CoV-2 drugs that were reported previously (Table 1 and Table S1). Together, our results demonstrate a comprehensive set of 91 approved drugs and 49 investigational drugs with anti-SARS-CoV-2 activity that can be considered for design of new clinical trials, especially drug combination therapies, to increase and improve treatment options for COVID-19.

## Discussion

In contrast to the other reported drug repurposing screens for SARS-CoV-2 using a single drug concentration in the primary screens (Dittmar et al., 2020; Ellinger et al., 2020; Jeon et al., 2020; Riva et al., 2020; Touret et al., 2020; Weston et al., 2020), we have used a quantitative HTS (qHTS, concentration-response) method (Inglese et al., 2006) where four compound concentrations were used in the primary screen instead of a single compound concentration. We also assessed the cytotoxicity of each compound against Vero E6 cells (without virus infection) in parallel with the SARS-CoV-2 CPE screening. The concentration-response for each compound used in the primary screen can improve identification of positive hits, especially these compounds with biphasic actions (bell-shaped curves) or screening errors. In addition, we have more inclusive compound collections with drugs approved by regulatory agencies outside of USA, such as Canada, Europe and Japan, that were not previously screened in SARS-CoV-2 assays. We also screened a set of investigational drugs that have human clinical data for drug properties such as the mechanism(s) of action, pharmacokinetics, and drug toxicity, which could be leveraged to speed up drug development. The other bioactive compounds screened have drug targets and mechanisms of action that may be useful for further studies of disease pathophysiology and for potential drug development.

We identified 319 compounds with activity against SARS-CoV-2 CPE from a qHTS of 8,810 unique compounds. Among the top 56 hits identified with <10 μM EC_50_ values and >80% efficacies, the anti-SARS-CoV-2 activity of 37 of them has not been reported elsewhere. Of these novel top hits, three were FDA approved drugs with novel anti-SARS-CoV-2 activity. Chlorprothixene is a dopamine receptor antagonist, a classic antipsychotic agent approved for treatment of schizophrenia (Schrijver et al., 2016). Methotrimeprazine, also named as levomepromazine, is another tricyclic antipsychotic agent approved for psychotic disorders including schizophrenia, and manic-depressive syndromes (Sivaraman et al., 2010). Both chlorprothixene and methotrimeprazine were previously found to inhibit the SARS-CoV replication with EC_50_s around 10 μM (Barnard et al., 2008). Piperacetazine is also an older tricyclic antipsychotic drug approved for treatment of schizophrenia (Eslami Shahrbabaki et al., 2018). The antiviral effect of piperacetazine was found previously that blocked the Ebola viral entry with the EC_50_ of 9.68 μM (Kouznetsova et al., 2019).

We also confirmed the anti-SARS-CoV-2 activity of 5 compounds that were reported as virtual screening hits but had yet to be confirmed experimentally, including methdilazine by an AI prediction algorithm (Grzybowski et al., 2020), GMC 2-113 by a virtual screen of RNA dependent RNA polymerase (RdRP) (Dwivedy et al., 2020), maprotiline by a main protease docking (Chauhan, 2020), deserpidine by a NPS-16 docking (Jiang et al., 2020), and flunarizine by a spike protein docking screen (Chernyshev, 2020). Our data supports the utility of these emerging technologies and the field of AI for advancing drug development.

For *in vitro* screens of antiviral compounds, molecular target (mechanism) based assays and phenotypic assays are two major approaches. Common targets are viral enzymes such as viral protease, DNA and RNA polymerases, reverse transcriptase, and integrase. Development of assays targeting viral enzymes rely on viral enzyme expression, purification, assay development, and validation (Shyr et al., 2020). Alternatively, phenotypic assays involving live virus infections are readily executed once the viruses are isolated from patients and viral replication in appropriate host cells are established. A common live virus infection assay is the measurement of CPE in virus infected host cells. There are two possibilities (fates) for the host cells after viral infection, including cytopathic infection (i.e. death of host cells) and persistent infection (Heaton, 2017). The CPE effect can be readily measured by the ATP content cell viability assay, which is robust and amenable for HTS. Due to the nature of the CPE assay, compounds that suppress CPE can act against any part of the virus infection cycle, including the binding of virus to host cell receptor, entry into host cells, virus replication, viral assembly/budding, and virus reinfection of adjacent cells.

It is worth briefly reflecting on the limitations of the drug repurposing assay approach. A number of small molecules of interest for treating COVID19 that are currently in clinical trials were not hits in our assay. For example, the TMPRSS2 inhibitors camostat and nafamstat are protease inhibitors approved in Japan for treating pancreatitis, and known to inhibit TMPRSS2 (Shrimp et al., 2020). While TMPRSS2 is reported to be a mediator of SARS-CoV-2 cell entry, Vero E6 cells do not express TMPRSS2, so this class of compound are not active in the Vero E6 assay. The drug efflux transporter P-glycoprotein (P-gp) can reduce cellular concentrations of test agents, and as a kidney epithelial cell line, Vero E6 cells likely expresses significant P-gp concentrations, which would reduce activity of P-gp substrates (Robey et al., 2018). Remdesivir itself is a substrate of Pgp (EMA, 2020), and is weaker against SARS-CoV-2 in assays using Vero E6 cells (EC_50_ > 1 μM) compared with Calu-3 or Huh7 cell lines (EC_50_ > 50 nM) (2020). These examples highlight the need for careful interpretation and critical follow-up studies after initial high-throughput screening analyses.

Importantly, the comprehensive primary screen datasets of this study for approved and investigational drugs, and mechanism-based bioactive compounds have been made publicly available in real-time on the NCATS OpenData Portal (https://opendata.ncats.nih.gov/covid19/index.html) (Brimacombe et al., 2020). These datasets provide a wealth of quality live-virus data that is freely available to the research community for future studies and data mining with the aim of offering a new medicine to treat COVID-19 patients quickly and safely (Huang et al., 2020; Zhu et al., 2020a).

## Acknowledgements

This work was supported by the Intramural Research Program of National Center for Advancing Translational Sciences, National Institutes of Health, Bethesda, MD, USA.

## Author contributions

PS, ZI, RTE prepared the assay ready plates. RB and LR conducted the CPE and cytotoxicity assays. PS, CK-T, KMW and SGM curated the compound libraries. CZC and MDH designed the experiments. CZC, BB, WZ wrote the manuscript. RH, MS, XH, HG, TZ performed data analysis and data uploads. All authors provided critical reading of the manuscript.

